# Ubiquitous predictive processing in the spectral domain of sensory cortex

**DOI:** 10.1101/2025.07.31.667946

**Authors:** Eli Sennesh, Jacob A. Westerberg, Jesse Spencer-Smith, Andre Bastos

## Abstract

The appearance at the anatomical level of a canonical laminar microcircuit suggests that each six-layer column of granular cortex may mediate a canonical computation. Hypotheses for such computations include predictive coding, predictive routing, efficient coding, and others. However, single-neuron recordings capture only the individual elements of the hypothesized laminar microcircuit, while local field potentials (LFPs) from a laminar probe offer insight into the broader population activity. Through the Allen Institute’s OpenScope Brain Observatory, data in mice performing a visual oddball task during multi-area laminar recording was used to test predictive processing hypotheses in the spectral domain. Histological labeling of the cortical laminae enabled a fine-grained examination of their roles in the task, and frequency bands capturing both feedforward and feedback effects were analyzed. ɣ-band local-field potential (LFP) oscillations conveyed feedforward prediction errors in lower sensory areas of cortex; ⍺/β-band oscillations weakened in unpredictable conditions compared to predictable ones; and θ-band oscillations additionally signalled slower, longer-scale temporal prediction errors. In combination with the previous findings, predictive routing explains these experiments where neither ubiquitous predictive coding nor feedforward adaptation can.

**Significance:** Cortical columns robustly signal perceptual features through the firing rates of spiking neurons. In accordance with this rate coding, predictive processing theories hypothesized that neuronal firing rates ubiquitously signal surprise. However, a recent large-scale study of spike rates did not support this conjecture. An alternate model, predictive routing, suggests that neuronal oscillations rather than spike rates could encode surprise. These neuronal oscillations, which can affect the timing but not rate of spiking, formed coherent ɣ rhythms which consistently signaled both simpler and more complex forms of surprise in mouse visual cortex. Together with the findings on spike-rates in the same experiment, our findings suggest that cortical circuits encode surprise in the rhythmic timing of spikes rather than in their rate.

## Main Text

The brain is continually bombarded with noisy, ambiguous sensory signals from which it must construct actionable percepts via an internal model. Oddball paradigms featuring repeated, or otherwise patterned, stimuli allow experimentation on the neural basis of this internal model. Responses to repeated stimuli decline with added repetition(*1–4*), and many measurement modalities that directly (e.g., the local field potential (LFP)) or indirectly (e.g. blood-oxygen level dependent (BOLD) responses(*5*)) derive from collective circuit activity also appear to reflect the predictability of stimuli(*6*). These observations have motivated the theoretical paradigm of *predictive processing*(*7–9*), founded upon early theoretical studies of predictive coding in the retina(*10*) and primary visual cortex(*11*).

Predictive coding (PC) hypothesizes that sensory perception consists of combining top-down and bottom-up signals in a canonical cortical microcircuit(*12*). PC models the top-down signals as “predictions” that serve to both explain away anticipated stimuli and to disambiguate uncertain stimuli, while hypothesizing that aspects of the stimulus not thus explained ascend the cortical hierarchy as bottom-up “prediction errors”. Experimentally, previous studies have used various auditory and visual oddball paradigms that first set up a predictable pattern, typically by means of stimulus or pattern repetition, and then violate it(*13*). Neuronal responses to the violations are interpreted as prediction errors.

The question of which neuronal circuits implement predictive processing in the brain remains open. Previous work has focused on decoding oddballs from evoked spiking rates(*6*, *14–17*). A recent large-scale neurophysiological survey(*17*) of visual cortex in both mice and monkeys reported on the same visual oddball paradigm used here, but confined its analyses to the rate-coded spiking domain. While this previous study did found that some prediction errors were detected in higher-order areas (LM, PM, and AM in mice) and fed back down, it did not test PC models’ hypotheses (*12*, *18–21*) in the spectral domain.

A growing body of work (*22–28*) has linked predictive processing to the oscillatory dynamics of LFPs. Presently, a controversy exists regarding which bands of cortical oscillations correspond to which theoretical constructs in predictive processing, specifically regarding the role of ɣ-band oscillations. Some authors(*24*, *29*) have characterized ɣ oscillations as synchronizing the superficial layers of neighboring cortical columns according to the spatial predictability of the stimuli driving them, while other authors(*25*, *27*, *30*) have associated θ and ɣ oscillations with prediction errors and ⍺/β oscillations with predictions. The latter accounts enjoy compatibility with findings(*22*, *31–34*) that ⍺/β and ɣ oscillations respectively mediate feedback and feedforward signaling. We describe this account of feedback and feedforward signaling, with ⍺/β exerting inhibitory control over ɣ in terms of the proposed general cortical mechanism of “predictive routing”(*27*). Still other authors(*35*) provided evidence that in a roving oddball paradigm, the prediction error signal appears in the aperiodic component of LFP spectrograms. Under this perspective, spectral changes resulting in increases in ɣ and decreases in ⍺/β (or vice-versa) are better captured by aperiodic shifts rather than changes to specific neuronal oscillations(*36*).

The contradictions between the above accounts require careful experiments that disentangle the different possible neuronal mechanisms purported to generate prediction-error signals, as well as analysis that takes into account the distinctions argued in the literature. This paper tests core tenets of the (hierarchical) predictive routing model using Multi-Area, high-Density, Laminar-resolved Neurophysiological (MaDeLaNe) recordings of local field potentials in mice; these mice underwent a large number of pre-recording repetitions of the “xxxy” pattern displayed in the Global/Local Oddball paradigm (*13*) to ensure that they would form perceptual expectations about the sequence and attenuate any novelty responses to the sequence. We analyzed data from six visual cortical areas, from the primary visual cortex (V1) to AM(*37*) at the top of the mouse visual hierarchy. After examining oscillatory activity in the spectral space, we also wanted to know whether the oscillatory bands analyzed routed information in the feedforward or feedback directions within the cortical column. Previous work in macaque areas V1(*27*) has identified a laminar pattern of ɣ current sinks feeding from cortical layer 4 to more superficial and deep layers, laminar pattern of activation classically associated with feedforward processing(*12*, *38*). Previous work was also found that ⍺ sinks originate in superficial and deep layers and move towards layer 4, a pattern classically associated with feedback processing. We used these same distinctions to test for laminar feedforward/feedback flow in the ɣ, ⍺/β and θ frequency ranges during oddball processing.

These techniques enabled us to test the following core hypotheses of predictive routing in both local and global oddballs, as well as random deviants. **H1**: Predictive routing suggests that feedforward ɣ increases in the processing of unpredicted stimuli, while feedback ⍺/β serves to selectively suppress the processing of predicted stimuli. This results in a “push-pull” dynamic between ⍺/β and ɣ. H1 further suggests a role for θ oscillations in prediction error processing as well. **H2**: Counter to H1, the “ɣ as prediction” model suggests that ɣ synchronization reflects the confirmation of predictions by an unsurprising stimulus, as suggested by previous physiology work in the spatial(*24*) domain and modeling work across spatial(*29*) and temporal(*26*) domains. **H3**: Predictability vs unpredictability tilts or shifts the aperiodic component of the spectral response, rather than affecting oscillations within specific frequency ranges.

We tested H1-H3 by submitting the data to analysis during three different oddball types, which we term “deviance detection”, “global oddball”, and “stimulus-specific adaptation”. Importantly, which stimulus-specific adaptation tests how the brain responds to a change of stimulus, deviance detection and global oddball contrasts both test for effects of prediction while controlling for the effects of short-term adaptation. Consistent with **H1**, contrasting these three different types of oddballs against their corresponding controls demonstrated a clear, significant increase in ɣ oscillations between 30-60 Hz, although the timing, laminar origin, and center frequency of these responses differed across oddball types. Consistent with **H1** but inconsistent with **H2**, global oddballs increased ɣ oscillations, showing that violation of predictions nonetheless drove an increase in ɣ power. Partially consistent with **H3**, each oddball induced a significant spectral shift in at least one area, but only in the first 150 milliseconds of spectral response.

## Results

Animals were first habituated extensively to local oddball sequences in two cohorts over several days (Figure 1a): one to an “xxxy” sequence and the other to a “yyyx” sequence. The ‘a’ and ‘b’ stimuli within a sequence denote drifting gratings at 135 or 45 degrees from horizontal; in the rest of the paper, ‘x’ will refer to the angle of stimulus used in the first three stimuli of the habituation sequence and ‘y’ to the angle of stimulus used in the last stimulus of the habituation sequence. Lower-case letters will denote fully predictable sequence elements (probability 𝑃 = 100%), while capital letters will denote randomized or oddball sequence elements (with the respective probability of occurrence defined in the text where relevant). The paper will proceed to describe the habituation sequence as “xxxy”. The global oddball sequence “xxxX” sequence did not occur during training, nor during the initial 50 habituation trials (Figure 1b) of the recording session. Only the main block contained the “xxxX” sequence in 20% (randomized) of its trials (Figure 1b).

**Figure 1.**
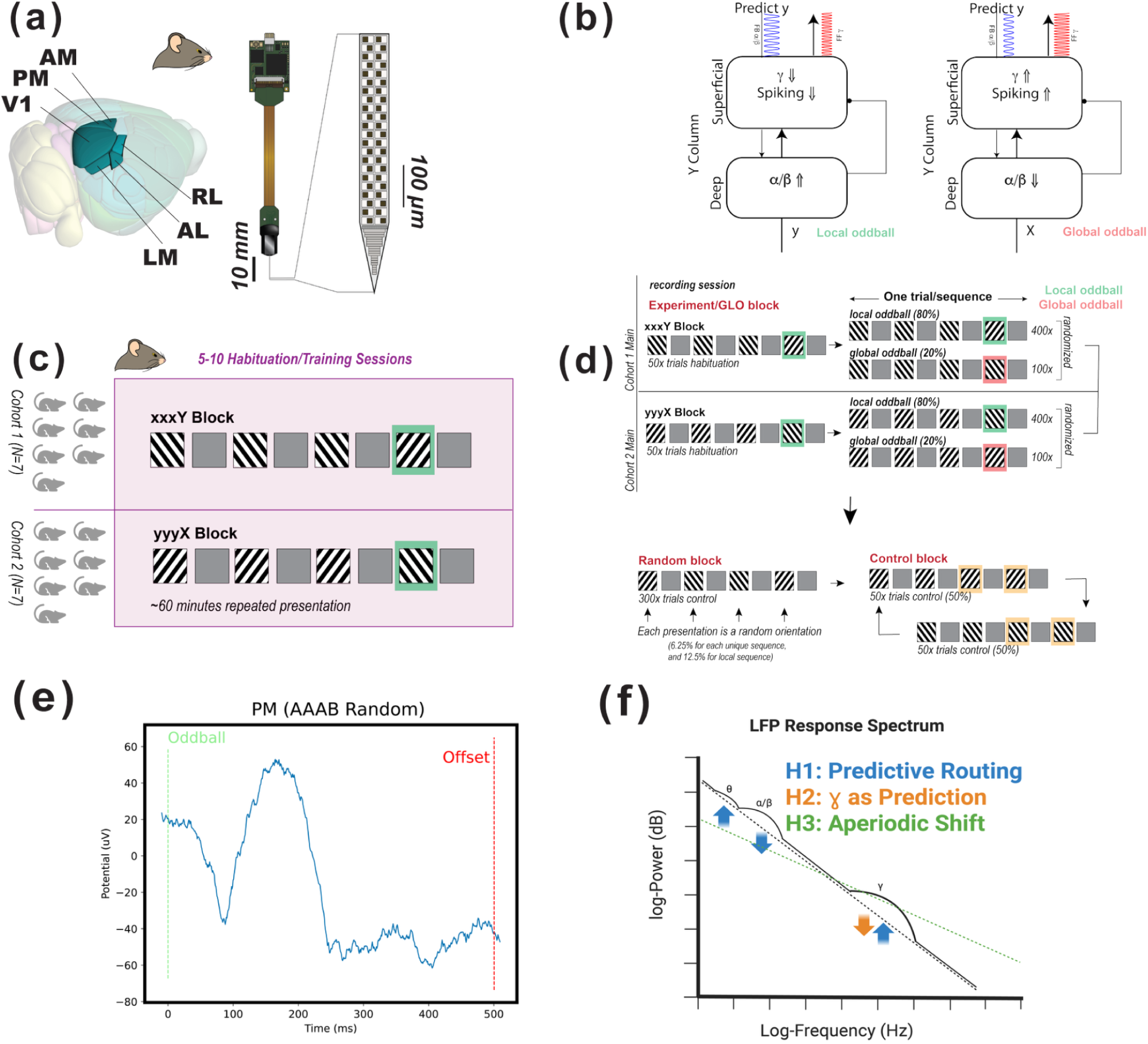
Local/global oddball paradigm, experimental setup, and predictive routing. **(a)** Neuropixels were introduced into 6 visual cortical regions in all mice (V1, LM, RL, AL, PM, AM). **(b)** How predictive routing models local and global oddballs: in the local oddball, a feedback ⍺/β oscillation predicting stimulus ‘x’ will fail to inhibit cortical columns responsive to stimulus ‘y’, resulting in uninhibited ɣ oscillations and spiking upon the onset of an unexpected ‘Y’, while in the global oddball, a feedback ⍺/β oscillation predicting stimulus ‘y’ after “xxx” will fail to inhibit cortical columns responsive to stimulus ‘x’, resulting in uninhibited ɣ oscillations and spiking upon the onset of the unexpected repetition ‘X’. **(c)** N=14 animals were habituated to the local oddball sequences (“xxxY” in Cohort 1 of 7 animals, “yyyX” in Cohort 2 of 7 animals) before recording; stimuli were presented for 500 ms with a 500-531 ms inter-stimulus interval and 1 s inter-trial intervals. **(d)** Visual stimuli were presented in 4-stimulus sequences consisting of two (2) oriented, drifting bar gratings (symbolized as ‘x’ or ‘y’), each displayed full-screen. After the main block of recording in which local and global oddballs were presented, the random control block consisted of randomized individual stimuli presented in sequences of four, while the repetition control block consisted of sequences of four repetitions of the same stimulus. **(e)** Local field potential in the middle channel of L4 over the course of the ‘y’ stimulus in the random deviant condition in the higher-order visual area PM; the plot shows oscillatory activity before, during, and after the stimulus-driven negative deflection in the local field potential. **(f)** Schematic of how surprise affects spectral responses according to hypotheses **H1** (predictive routing), **H2** (ɣ encoding prediction), and **H3** (aperiodic shifts). Dotted lines show the aperiodic component of the spectral response, while solid lines show the reconstructed spectrum that includes oscillatory peaks. **H1** (blue) predicts that surprise increases θ power, decreases ⍺/β power, and increases ɣ power; **H2** (orange) predicts that surprise decreases ɣ power while confirmation increases it; and **H3** (green) predicts that surprise shifts the aperiodic component of the spectrum. Created in BioRender. Sennesh, E. (2025) https://BioRender.com/ogllv24.

To investigate the neural signaling underlying prediction of predictable sequences and processing of unpredicted stimuli, we compared responses in the main block with those in two control blocks (Figure 1d): a random block (in which “XXXX” had probability 𝑃 = 6. 25%, the same as “XYXY”) and a fully predictable block (in which sequences were predictable from their first stimulus, so that “xxxx” had 𝑃 = 50% and “yyyy” had 𝑃 = 50%). To analyze oddball responses, we compared spectral responses to the fourth stimulus in the sequence (denoted P4) across blocks. First, to examine modulation of responses by probabilistic context, we contrasted P4 in the randomized control block (‘Y’ in “XXXY”) with P4 in the fully deterministic habituation block at the beginning of recording (‘y’ in “xxxy”); we refer to this contrast as “deviance detection”. Second, to examine the response to prediction error without a release from stimulus-specific adaptation, we contrasted P4 in the global oddball (the ‘X’ in “xxxX”) against P4 in the deterministic control block (the last ‘x’ in “xxxx”). We refer to this contrast as the “global oddball”. Third, to examine the spectral effects of release from stimulus-specific adaptation, we contrasted P4 in the local oddball (the ‘Y’ in “xxxY”) against P4 in the deterministic control block (the last ‘y’ in “yyyy”). We refer to this contrast as “stimulus specific adaptation”.

After the spectral decomposition of the LFPs was performed, all contrasts consisted of a channel-time-frequency (three-dimensional) nonparametric, cluster-based statistical test(*39*) with a family-wise error rate ⍺ = 0. 01; the entire paper uses this significance level and testing procedure for its statistical tests. The tests thus resulted in three-dimensional significance masks for the difference in trialwise means of spectrograms. All reported results passed significance at this threshold (see Methods below for further details). We also performed the aperiodic vs oscillatory split analysis specifically for the first 150 ms of the spectral response, a feedforward sweep before feedback signaling comes online during stimulus presentation(*40*, *41*); Supplementary Figure 3 shows the results of this analysis and will be referred to throughout the presentation of results.

### Narrowband ɣ oscillations reflect deviance detection

The Global-Local Oddball paradigm(*13*) contains the standard oddball sequence in two different probabilistic contexts: the habituation block’s “xxxy” sequences (𝑃 = 100%) and the random control block’s fully randomized “XXXY” sequences (𝑃 = 15%), the latter following the abolition of the habituation and therefore abolition of the highly predictable nature of “xxxY”. Combining these for our deviance detection contrast enabled testing **H1**, **H2**, and **H3** via the effects of context upon responses to the standardized oddball sequence.

**H1** hypothesizes that unpredicted but attended stimuli evoke feedforward ɣ oscillations, while suppressing feedback ⍺/β oscillations, and that probabilistic uncertainty, not just stimulus identity, modulates the power of these oscillatory responses, while **H2** hypothesizes that ɣ oscillations reflect confirmation of predictions (and thus, certainty rather than uncertainty). **H3** hypothesizes that predictability modulation should affect the aperiodic component of the spectral response, rather than the oscillatory component. Our experimental design enables testing these hypotheses with the three contrasts defined above. We note that a possibility is that ɣ, ⍺/β, and aperiodic components may behave differently for each contrast, and alternatively, these signals may also behave consistently across the contrasts if they are canonically related to stimulus (un)predictability.

We begin with the deviance detection contrast in area V1. All analyses have been performed after removing the FOOOF-computed aperiodic component as a baseline, and are therefore more interpretable as genuine oscillatory activity. Figure 2 shows the oscillatory spectral responses to the fully randomized final ‘Y’ presentation of the sequence “XXXY” (𝑃 = 15%) in contrast to those to the fully deterministic ‘y’ presentation of the sequence “xxxy” (𝑃 = 100%) during habituation. Figures 2(a) and 2(b) show the spectrograms relative to the FOOOF-computed aperiodic component, while Figure 2(c) shows the spectrogram of the deviance detection contrast (see Methods for our channel pooling strategy of across trials and animals). We observed a characteristic decrease in ⍺/β power (1-2 dB) just after stimulus onset, followed by ɣ oscillations (1-2 dB) that began at the 40-65 Hz frequency band and descended down towards the 35-55 Hz band over time. Since the cluster masks produced by our statistical tests are three-dimensional, Figures 2(d-f) display the laminar activation patterns corresponding to the full ɣ (30-90 Hz), ⍺/β (10-30 Hz), and θ (2-10 Hz) bands. Figure 2(d) shows that ɣ oscillations are strongest in superficial layers 1-4 (around 2 dB increase in oddball-related power); after approximately 100 ms a weak (<1 dB) but significant loss of ɣ power occurs in laminar L5/6 while a gain in synchronization (1-2 dB) comes to dominate laminar L2/3. Figure 2(e) shows that increases in ⍺/β power (1-1.5 dB in power) occurs throughout stimulus presentation in L1 of V1, though it is interrupted by a powerful (2 dB) desynchronization beginning in laminar L2/3 and spreading down into L4 and L5/6. Figure 2(f) shows that θ synchronization occurs only weakly (< 1 dB) in L1 of V1 during deviance detection. To determine the laminar origin of current sinks and sources of these LFP oscillations, and thus whether they index feedforward or feedback captivity, we analyzed the current source density relative to the troughs of these oscillations (see Methods for details). Figures 2(g-i) show the trough-locked current-source density profiles of the ɣ, ⍺/β, and θ oscillations.We observed a feedforward pattern of activation in ɣ oscillations, whereby sinks originated in the middle layers and spread out to superficial L2/3, followed by later sinks in L5/6. There was a characteristic laminar feedback pattern in ⍺/β oscillations, which originated in the most superficial contacts and spread to the middle layers. θ oscillations displayed a mixed pattern consistent with aspects of both a feedback and feedforward pattern.

**Figure 2.**
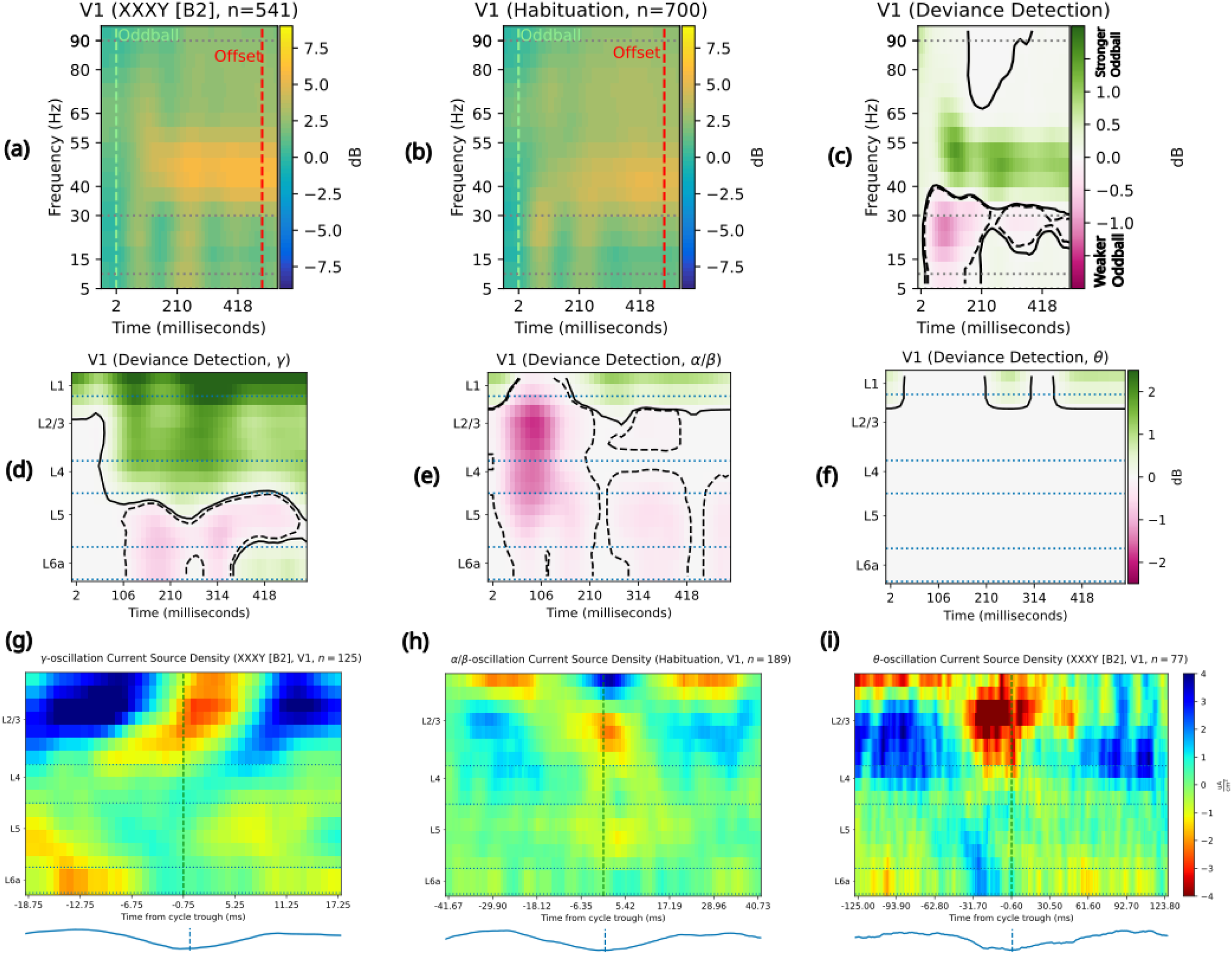
Probabilistically deviant stimuli induce a “push-pull” between. ⍺**/β- and ɣ oscillations.** The top row shows the two conditions contrasted to examine deviance detection; the middle row shows the laminar signatures of the contrast results broken out by frequency band; and the bottom row shows the laminar CSD responses characteristic of each frequency band in these conditions. Solid contour lines enclose the area of a synchronization/power increase, while dotted contour lines enclose the area of a desynchronization/power decrease. **(a)** Spectral responses to a fully randomized “XXXY” (n=541 trials) conform to stereotyped “oddball” responses, with stimulus onset leading to an increase (2.5-5 dB) and then decrease (0-2.5 dB) in ⍺/β power over baseline, then to a long-lasting increase (2.5-5 dB) in ɣ (40-55 Hz) synchronization. **(b)** Spectral responses to a fully predictable, habituated “xxxy” (n=700 trials) show a fast increase (>5 dB) over baseline and then return to baseline of ⍺/β power, followed by a long-lasting increase (2.5-5 dB) in ɣ (40 rising to 40-55 Hz over time) synchronization. **(c)** Detection of probabilistic deviants, via the contrast between “XXXY” and “xxxy”, induces a 1 dB decrease in ⍺/β power and a 1-2 dB ɣ power increase. **(d)** Deviance-induced ɣ (30-90 Hz) synchronization occurs earliest and most strongly at the boundary between L1 and L2/3, remaining strong in L1 while spreading down into L2/3; a 0-1 dB loss of ɣ power occurs in L5/6 100ms into deviance detection. **(e)** Deviance-induced ⍺/β (10-30 Hz) activity starts with a fast-onset increase in synchronization in L1, followed by ⍺/β desynchronization (1.5-2 dB) in L/3 and down into L5, followed by a slow decrease in ⍺/β power (0.5 dB) in L5/6 and increase in ⍺/β power (1-1.5 dB) in L1. **(f)** Deviance detection induces a weak (0.5-1 dB), late increase in θ (2-10 Hz) power. **(g)** ɣ oscillations in P4 of the fully randomized “XXXY” sequence display a canonical feedforward pattern of laminar activation, beginning in L4, then flowing out to L/3 and later into L5/6, over n=125 complete oscillatory cycles re-epoched from 541 trials. Oscillatory LFP trace shown below for illustrative purposes only. **(h)** ⍺/β oscillations in P4 of the fully habituated “xxxy” sequence display a weak but canonical feedback pattern of laminar activation, beginning in L1 and flowing down to L2/3, then down through L4 to meet a rising flow of activation in L5. CSD was calculated over the average of n=189 complete oscillatory windows over 700 trials. Oscillatory LFP trace shown below for illustrative purposes only. **(i)** θ oscillations in P4 of the fully randomized “XXXY” sequence display a canonically feedback pattern of laminar activation, beginning in L1 and L6 before converging up through L4 to L2/3, followed by a feedforward pattern to L1 and L5/6 from L2/3. CSD was calculated over the average of n=77 complete oscillatory cycles out of 541 trials. Oscillatory LFP trace shown below for illustrative purposes only.

To ensure that apparent deviance detection results in fact reflect neuronal oscillations, we also separated the spectrum within each condition for each animal into its aperiodic and oscillatory components via FOOOF; Supplemental Figures 1 and 2 show these results. Deviance detection displayed no statistically significant difference between the aperiodic components of the spectral response to its oddball and its control stimulus in any of the recorded cortical areas. After removal of the aperiodic component, however, only V1 (𝑝 < 0. 05) and LM (𝑝 < 0. 01) displayed significant increases in both the power and peak frequency of ɣ oscillations, derived from a permutation-based *t*-test (with 𝑛 = 1000 permutations) for difference of means between conditions on per-subject, channel- and trial-averaged frequency spectra. Supplementary Figure 3 shows that deviance detection induced a significant difference in the aperiodic spectral response of the feedforward sweep of 150 ms only in LM.

These findings show consistency with hypotheses **H1** and inconsistency with **H2**. Consistent with predictive routing (**H1**), ɣ power increased, ⍺/β power decreased, and θ power increased to an unpredictable stimulus vs a predictable one. Inconsistent with **H2**, the random deviant “XXXY” exhibited higher, rather than lower, ɣ power than the fully predictable “xxxy” in habituation. **H3** concerns the aperiodic component of the spectral response, so combined with the results in Supplemental Figures 1, 2, and 5 these results are mostly inconsistent with **H3**, excepting the difference in LM in the aperiodic component.

### ɣ oscillations responded to global oddballs

The Global-Local Oddball paradigm(*13*) contains two conditions in which stimulus repetition occurs without a release from neuronal adaptation: during the main block, the global oddball “xxxX” occurred with 𝑃 = 20%, while during the repetition control block, “xxxx” occurred during half the trials and its final stimulus ‘x’ occurred with 𝑃 = 100% given the first stimulus. This global oddball contrast enabled testing **H1**, **H2**, and **H3** without the effects of any release from stimulus-specific adaptation.

Figure 3 shows the oscillatory spectral responses, measured in decibels of difference across conditions, to the global oddball “xxxX”, in contrast to those to the deterministic control sequence “xxxx”. Figure 3(a) shows the response to the final ‘X’, while Figure 3(b) shows the response to a predictable ‘x’ in the control condition. Both responses consist of an increase in ⍺/β power over baseline at the onset of the stimulus, alongside an increase in ɣ (30-90 Hz) power in 40-55 Hz and 80-90 Hz bands that lasts throughout stimulus presentation. However, in the global oddball condition the ⍺/β band quickly returns to baseline, while the ɣ oscillations increase more in power (2 dB vs <2 dB) compared to the control block. Figure 3(c) shows the statistically significant contrast between the global oddball ‘X’ and stimulus control ‘x’, derived from a channel-time-frequency nonparametric statistical test with family-wise error rate ⍺ = 0. 01 in which spectrogram data was averaged over channels; the contrast response consists of a fast increase (>=0.3 dB) in high-ɣ (55-90 Hz) synchronization, which trails off into sustained increases (<= 0.2 dB) in 40-55 Hz ɣ power, alongside an immediate rise (0.2-0.3 dB) in θ power and a fall (<0.2 dB) in ⍺/β power over time. We interpret the low-frequency synchronization shown in Figure 3(c) as θ synchronization in light of a further spectral analysis with higher frequency resolution, shown in Supplemental Figure 2, finding oscillatory peaks at 5-9 Hz, within the <10 Hz θ band. Figures 3(d), 3(e), and 3(f) each display one frequency band’s laminar profile over time. Figure 3(d) shows that an immediate increase in ɣ (0.4-0.6 dB) power occurred in the supragranular laminar layers and grew to >= 0.6 dB over the course of global oddball presentation. Figure 3(e) shows that an immediate ⍺/β loss of power (0.75-1 dB) took place in upper supragranular layers, while a weak increase in ⍺/β power spread from infragranular layers upwards after approximately 120 ms. Figure 3(f) shows that within less than 100 ms after oddball onset, an increase in θ power (< 0.5 dB) begins in infragranular layers before intensifying to 1.0-1.5 dB as it spreads throughout V1. Figures 3(g-i) show the current source density profiles of the ɣ, ⍺/β, and θ oscillations, time-locked to the oscillation troughs; these demonstrate a pattern of feedforward activation in ɣ oscillations, a feedback pattern in ⍺/β oscillations, and a pattern of feedback (L5/6 to L4) followed by feedforward (L4 to L2/3 and L5/6) activation in θ oscillations.

**Figure 3.**
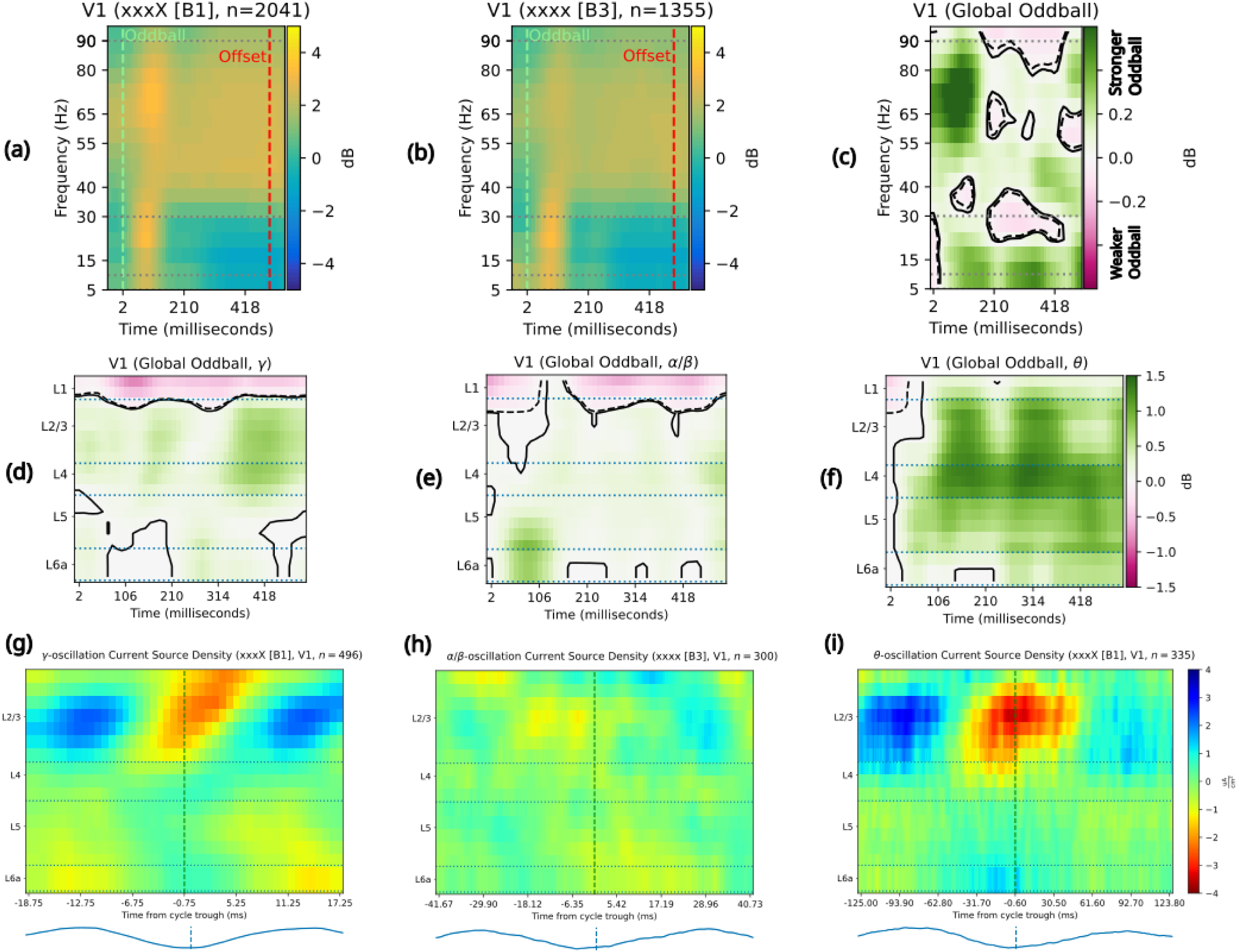
Unpredictable repetitions induce ɣ and θ oscillations. The top row shows the two conditions contrasted to examine global oddball detection; the middle row shows the laminar signatures of the contrast results broken out by frequency band; and the bottom row shows the laminar CSD responses characteristic of each frequency band in these conditions. Solid contour lines enclose the area of a synchronization/power increase, while dotted contour lines enclose the area of a desynchronization/power decrease. **(a)** Spectral responses to P4 in an unpredicted (20% probability, n=2041 trials) repetition “xxxX” show a fast increase (2-4 dB) in ⍺/β (2-4 dB) and increase (<2 dB) in high-ɣ power, followed by a return to baseline in the ⍺/β band, a <2 dB increase in θ synchronization, and long-lasting <2 dB increases in low-ɣ (40-55 Hz) and high-ɣ (80-90 Hz) synchronization. **(b)** Spectral responses to P4 in a fully predictable (n=1355 trials) repetition sequence “xxxx” show a fast increase (4-6 dB) in ⍺/β power and smaller (<2 dB) ɣ power increase over baseline; both bands of oscillations continued throughout the first 200ms. **(c)** Global oddball detection, via the ⍺ = 0. 01 statistical contrast between “xxxX” and “xxxX”, induces a 0.2 dB decrease in ⍺/β power after 200ms, alongside 0.4-0.5 dB increases in θ and ɣ power. **(d)** ɣ power increases by 0.2-0.4 dB in L5/6, L4, and L2/3 immediately upon onset of the global oddball, persisting and even strengthening to 0.6 dB by the end of stimulus presentation, while ɣ power decreased by 0.4-0.6 in L1. **(e)** ⍺/β power increase begins (0.5-1 dB) in L5/6 <100ms after global oddball onset, while ⍺/β power decreases (0.5-0.75 dB) in L1. The increase in infragranular layers spreads out to L4 and then to L2/3 while weakening to 0.25-0.5 dB. **(f)** θ oscillations begin in L5/6 with a 0.5dB increase just after global oddball onset before spreading and strengthening (1-1.5 dB) throughout the laminar column. **(g)** ɣ oscillations induced by global oddballs show a canonically feedforward laminar activation pattern, starting in L4 at phase 0 and spreading out towards L2/3 and L5/6 as the phase cycle continues. CSD was calculated over the average of n=496 complete oscillatory windows over 2041 trials. **(h)** ⍺/β show a weakly feedback-driven laminar activation pattern, starting in L1 before continuing down to L2/3 and L4, with later contributions coming from L5/6. CSD was calculated over the average of n=300 complete oscillatory windows over 1355 trials. **(i)** θ oscillations show a feedforward laminar activation pattern under global oddballs, with activation beginning in L5 and L4 before spreading to L2/3 as it strengthens with the phase cycle. CSD was calculated over the average of n=335 complete oscillatory windows over 2041 trials.

Ensuring that apparent global oddball results in fact reflect neuronal oscillations, the aperiodic component of the per-animal, per-condition spectra showed no significant differences between the last response in “xxxX” and “xxxx” sequences (Supplementary Figure 1), although significant spectral tilt does appear fast feedforward sweep in the first 150 ms (Supplementary Figure 3) in V1, AL, and PM. In fact, the primary significant differences driven by the global oddball were in the more predictable condition “xxxx” showing greater ⍺/β than the global oddball “xxxX” (Supplementary Figure 2) in V1, LM, RL, AL, and PM. We therefore interpret our global oddball results in terms of both spectral tilt and oscillation dynamics.

These findings are consistent with **H1**, inconsistent with **H2**, and partially consistent with **H3**. The pattern of combined ɣ and θ synchronization with ⍺/β desynchronization, driven by an unpredictable stimulus, fits **H1**. Both the global oddball and control conditions feature identical stimulus sequences, but the ɣ power increase for the surprising (𝑃 = 20%) global oddball rather than predictable (𝑃 = 100%) control is inconsistent with **H2**. **H3** hypothesized that unpredictability should affect the aperiodic component of the spectrum rather than the oscillatory one; the findings in Figure 3 combine with those in Supplementary Figure 3 to show consistency with **H3** for the fast feedforward sweep of 0-150 ms post-onset, while combining with those in Supplementary Figure 1 to falsify **H3** for the later 150-500 ms period.

### Stimulus change induces ɣ vs ***⍺***/β push-pull

**H1** hypothesizes that classic, “local” oddballs should induce a push-pull dynamic between feedforward ɣ and θ synchronization and ⍺/β desynchronization**. H2** suggests that ɣ synchronization should reflect the confirmation of predictions by the stimulus. **H3** suggests that predictability or unpredictability should shift the aperiodic component of the spectral response rather than the oscillatory one. The Global-Local Oddball paradigm (*13*) enables testing for “local” oddballs (in which repetition creates an expectation which is violated via a change of stimulus content) for the effects of a release from stimulus-specific neuronal adaptation. This test consists of contrasting the oscillatory responses to a “local” oddball sequence “xxxY”, in which a release from adaptation would occur with stimulus change, with those to fully deterministic sequences having the stimulus content “yyyy”, in which neuronal adaptation would continue throughout all four presentations (see Figure 1(d) for a full description of sequence types).

Figure 4(a-c) shows the oscillatory spectral responses, measured in decibels above the aperiodic baseline, to the mostly predictable (𝑃 = 80%) final ‘Y’ presentation of the sequence “xxxY” (Figure 4(a)) in contrast to those of the fully deterministic (𝑃 = 100%) ‘y’ presentation of the sequence “yyyy” (Figure 4(b)). The contrast in Figure 4(c) shows non-frequency-specific desynchronization (0.5-1.5 dB) at the onset of the local oddball, followed by enhanced ɣ power (1-1.5 dB) throughout the stimulus presentation. Figures 4(d-f) each show the laminar profile of significant contrasts in oscillatory power within the ɣ, ⍺/β, and θ frequency bands. This confirms that the desynchronization is present in all frequency ranges and layers.

**Figure 4.**
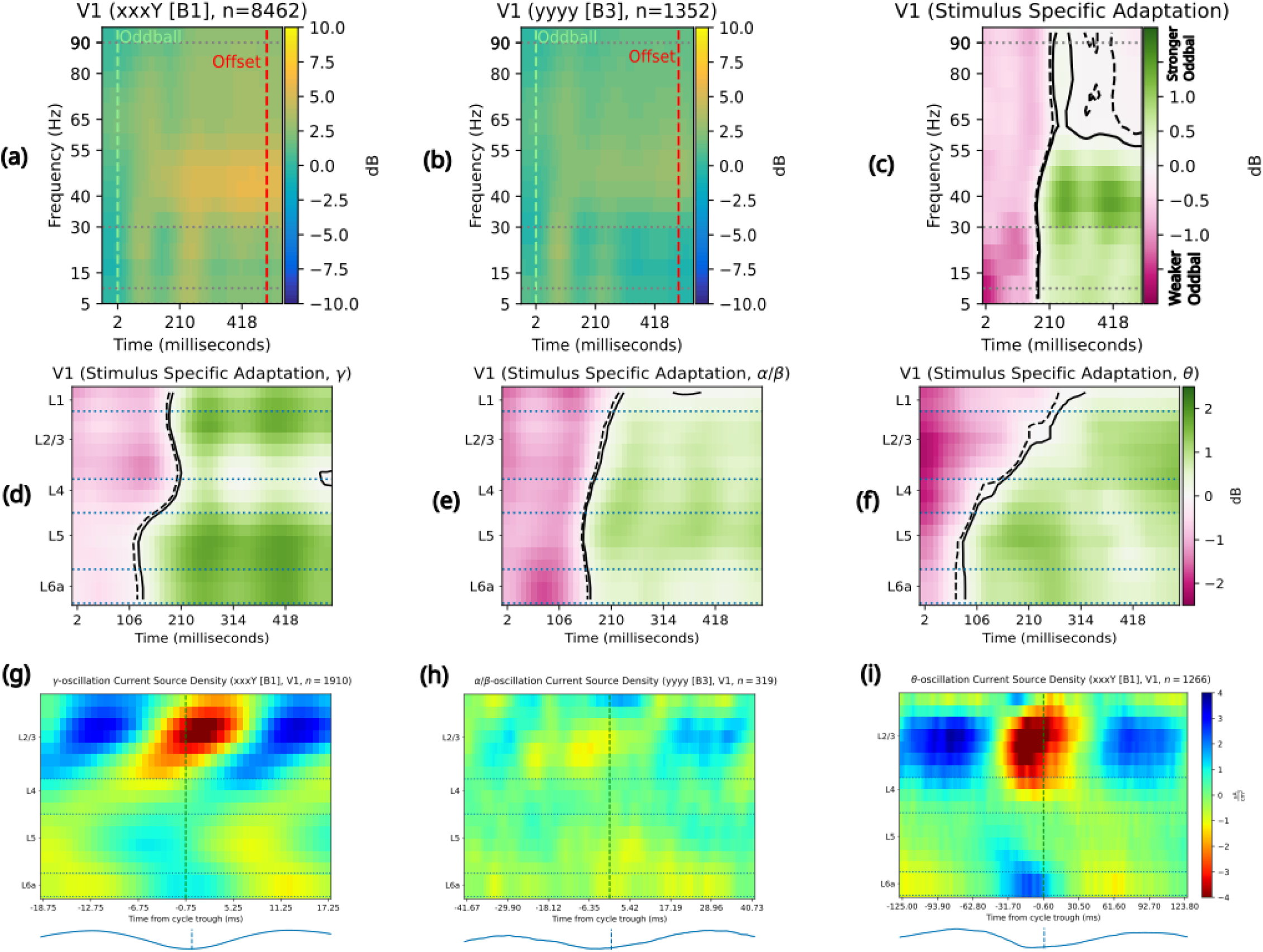
Release from stimulus-specific adaptation induces a “push-pull” between. ⍺**/β- and ɣ oscillations.** The top row shows the two conditions contrasted to examine release from stimulus-specific adaptation; the middle row shows the laminar signatures of the contrast results broken out by frequency band; and the bottom row shows the laminar CSD responses characteristic of each frequency band in these conditions. Solid contour lines enclose the area of a synchronization/power increase, while dotted contour lines enclose the area of a desynchronization/power decrease. **(a)** Spectral responses to P4 in a relatively predictable (80% probability) “xxxY” sequence conform to stereotyped “oddball” responses, with stimulus onset leading to an increase (2.5-5 dB) in ⍺/β power (followed by a return to baseline and then another, more sustained increase), accompanied by a fast-onset and then long-lasting increase (2.5-5 dB) in ɣ (40-55 Hz) and high-ɣ (80-90 Hz) synchronization**. (b)** Spectral responses to P4 in a fully predictable “yyyy” sequence show a fast increase (2-4 dB) in ⍺/β power after stimulus onset, followed by a smaller (<2 dB) increases in ɣ (40-55 Hz) and high-ɣ (80-90 Hz) synchronization. **(c)** Mostly-predictable release from adaptation, via the ⍺ = 0. 01 statistical contrast between “xxxY” and “yyyy” sequences, induces a 1-2 dB decrease in ⍺/β power at stimulus onset, followed by a 1-2 dB ɣ power increase and a roughly 1 dB increase in θ and ⍺/β power. **(d)** Release from stimulus-specific adaptation induces a fast, broad loss (<1 dB) of ɣ power, followed after 150ms by a 1 dB ɣ power increase from baseline that begins in L5/6 and spreads through L4 to L2/3. **(e)** Release from stimulus-specific adaptation induces a fast, broad loss (1-2 dB) of ⍺/β power, followed by a 1-2 dB increase from baseline in ⍺/β power that starts in L5/6 and spreads into L4 and L2/3 while never arriving to L1. **(f)** Release from stimulus-specific adaptation induces a fast, broad loss (1.5-3 dB) of θ synchronization that lasts longer in L2/3 than elsewhere, while after about 100ms a 1 dB increase in θ synchronization begins in L5/6 and spreads into L4 and L2/3 without ever arriving to L1. **(g)** ɣ oscillations in P4 of a mostly predictable “xxxY” sequence shows a canonical feedforward pattern of laminar activation, beginning in L4 and L5 at the end of one phase cycle and spreading out into L2/3 and L6 through the next cycle until the ɣ trough. CSD was calculated over the average of n=1910 complete oscillatory windows over 8462 trials. **(h)** ⍺/β oscillations in P4 of the fully predictable “yyyy” sequence show a feedback pattern of laminar activation, with current sinks spreading from L5/6 and L1 into L5, L4 and L2/3 before the end of the oscillation carries them into L1 and L6. CSD was calculated over the average of n=319 complete oscillatory windows over 1352 trials. **(i)** θ oscillations in P4 of the mostly predictable “xxxY” sequence show a mixed feedback and then feedforward pattern of laminar activity, with current sinks starting in L1 and L6 before activity in L2/3 at the θ trough spreads downward through L4 and L5 into L6 towards the cycle’s end. CSD was calculated over the average of n=1266 complete oscillatory windows over 8462 trials.

Additionally, Figure 4(d) shows a ɣ power increase (1-2 dB) that starts in L5/6 and spreads up through L4 into L2/3 and L1. Figure 4(e) shows that an increase in ⍺/β power (1 dB at its strongest, at approximately 150 ms) starting in L5 and spreading out to the rest of the cortical column. Figure 4(f) shows a θ synchronization (1-1.5 dB) beginning in L5/6 and spreading up into L4 and through to L2/3 by the time of local-oddball offset. Figures 4(g-i) show the laminar current-source density profiles of the ɣ, ⍺/β, and θ oscillations, time-locked to the oscillation troughs; these demonstrate a pattern of feedforward activation in ɣ oscillations (Figure 4(g)), a mixed feedforward and feedback pattern in ⍺/β oscillations (Figure 4(h)), and a pattern of canonical feedforward (L4 to L2/3 and L5/6) activation in θ oscillations (Figure 4(i)).

These findings appear most consistent with **H1**, partly inconsistent with **H2**, and partially consistent with **H3** (in light of the findings in Supplemental Figure 1 and Supplementary Figure 3). The push-pull effects between ⍺/β and ɣ oscillations conform to **H1**, but also motivated it. Partly inconsistent with **H2**, the change of stimulus involved in a local oddball increased ɣ power after the first 150 ms, rather than decreasing it; however, a ɣ increase in response to the local oddball (𝑃 = 80%) admits an explanation in terms of prediction confirmation via **H2**. Spectra for the full stimulus presentation displayed no significant effects of release from adaptation, inconsistent with **H3**, but spectra for the fast feedforward sweep in the first 150 ms, typically accompanied by high spiking, showed significance tilt in V1, AL, and PM, consistently with **H3** for that early sweep.

### Spectral power effects come from oscillations and aperiodic tilt

Spectra of neuronal local field potentials consist not only of oscillations but also of an aperiodic component whose strength tends to follow a 1/𝑓 law; without separating out this component from oscillations, false conclusions might be drawn (*42*). Supplementary Figures 1 and 2 show the results of applying FOOOF analysis to parameterize LFP spectra across the period of oddball processing (from 50 ms before oddball onset to 50 ms after oddball offset) within conditions and subjects into aperiodic and oscillatory components. Supplemental Figure 1 shows the aperiodic components calculated for oddball spectra calculated for each animal and condition, with the line showing the mean aperiodic component across animals and the error bands showing two standard errors of the mean across animals; no significant cross-condition differences appear in the aperiodic component. Supplementary Figure 2 shows that after removing the aperiodic component, oscillatory peaks in the power spectrum at θ and ɣ ranges exist in all areas and conditions. Interestingly, we observed a spectral shift in the ɣ frequency band most notably in the deviance detection contrast. We found that ɣ shifted on average 3 Hz frequency during the XXXY condition (mean ɣ peak in V1: 38 Hz with 99% confidence interval 35.2-40.5 Hz) compared to the same stimulus sequence during habituation trials (mean ɣ peak in V1: 35 Hz, 99% confidence interval 32.6-36.8). Supplementary Figure 3 shows that in the fast feedforward sweep of the first 150 ms after the stimulus presentation, V1, LM, AL, PM, and AM all exhibited a spectral tilt due to oddballs in at least one contrast, though no area showed a spectral shift in every contrast. Particularly in the case of the release from stimulus-specific adaptation, the spectral tilt in V1, PM, and AM fits with **H3**, despite the cross-condition contrasts in Figures 2-4 concerning the oscillatory activity after FOOOF removes the aperiodic spectrum.

Finally, we tested to what extent these effects in V1 could generalize to other visual cortical areas in mice. We analyzed ɣ-band oscillatory power in L2/3 across areas, and this demonstrated that all three types of oddballs show significant increases in ɣ power for oddball over control across multiple areas of the cortical hierarchy (with area RL in global oddballs being the one exception, Figure 5). Deviance detection and global oddball modulations were restricted to the early part of the hierarchy (areas V1, LM, RL, and AL for deviance detection and areas V1, LM, and RL for global oddball). Stimulus specific adaptation responses could be seen throughout the entire hierarchy of areas. These results are compatible with **H1**, because they show that ɣ acts as a signature of prediction error for multiple contrast types and in multiple areas. **H2** could be supported only by a single area and contrast, as area RL showed significant ɣ desynchronization during oddballs.

**Figure 5:**
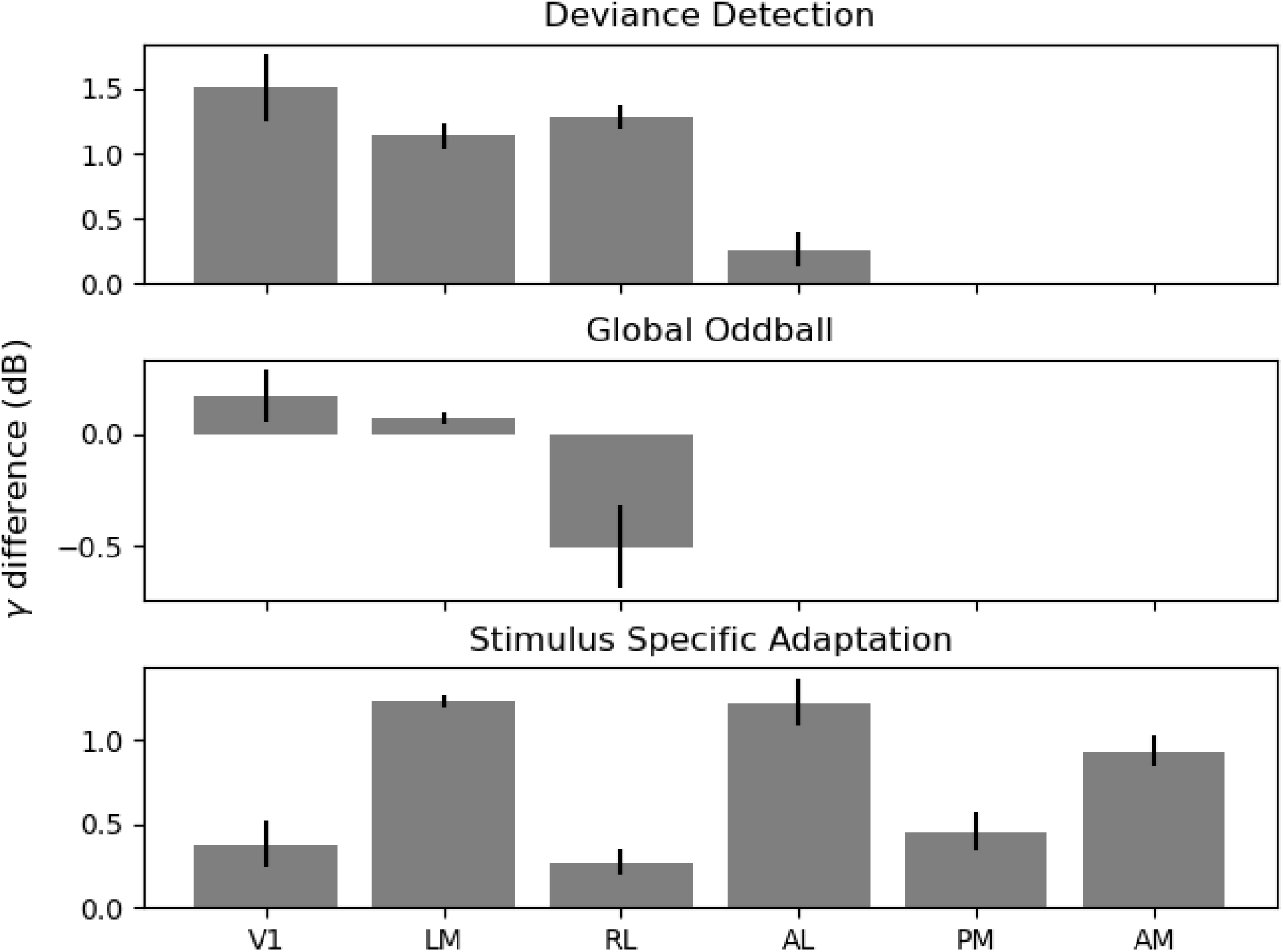
Early areas show greater, more significant differences in oscillatory ɣ power across deviance detection and global oddballs. The bar-plot above shows the time-average of ɣ power increases/decreases in the statistically significant clusters (⍺ = 0. 01) shown in Figures 2(c), 3(c), and 4(c) above bars show the mean (over channels) increase/decrease for each contrast while the error marks show two standard errors of the mean (over channels). Deviance detection induced a significant increase in oscillatory ɣ power of roughly 1.5 dB in V1, 1.1 dB in LM, 1.3 dB in RL, 0.3 dB in AL, and insignificant differences in PM and AM. Global oddballs induced smaller, but statistically significant (for channelwise statistics) increases in ɣ-band power of 0.2 dB in V1, 0.1 dB in LM, and -0.5 dB in RL, with no significant contrasts in AL, PM, and AM. Release from stimulus-specific adaptation generated significant increases in ɣ-band power across area, averaging to 0.4 dB in V1, 1.2 dB in LM, 0.3 dB in RL, 1.2 dB in AL, 0.4 dB in PM, and 0.9 dB in AM.

## Discussion

This paper aimed to test the hypotheses **H1-H3** about the oscillatory dynamics of visual cortical areas in the Global/Local Oddball paradigm(*13*) in mice. In every contrast of experimental conditions, unpredicted stimuli evoked increased ɣ power in superficial (L1-3) layers, with a feedforward current-source density pattern (✔ **H1**). All three oddball types also evoked ⍺/β desynchronization (✔ **H1**), although this occurred later and with a different laminar pattern in global oddballs than in deviance detection and release from adaptation. The deviance detection (randomization of stimulus change) and global oddball (unpredicted repetition) contrasts showed increased ɣ power with the violation, rather than confirmation, of predictions (✘ **H2**). The release from adaptation contrast showed fast (0-150 ms) decreases in ɣ power upon onset of a prediction-confirming stimulus (✘ **H2**), but this was followed by increased ɣ power after 150 ms (✔ **H2**). While response spectra across the full half-second stimulus presentation did not show an overall spectral tilt, those for the first 150 ms, during which feedforward spiking takes place, did show a spectral tilt in several areas and contrasts (✔ **H3**); differences in the late (150-500 ms) responses were confined to the per-trial oscillatory spectra and their dynamics.

Our results suggest that the feedback role of ⍺/β oscillations (see Figures 2(h), 3(h), and 4(h)) likely consistently involves layers L1/6. Since L1/6 are already responsible for delivering feedback from a higher cortical area in canonical microcircuit models(*43*), this revision mainly considers the laminar profile of ⍺/β oscillations rather than the participation of ⍺/β oscillations in feedback processing. The predictive routing model(*44*) had previously considered ⍺/β oscillations to come from L5/6 rather than L1/6, so ultimately our results suggest revising this to include a participation of both superficial and deep layers in ⍺/β generation(*45*, *46*). This participation of both the supragranular (L1) and infragranular (L6) targets of corticocortical feedback in ⍺/β generation may be one reason why mice lack the “spectrolaminar motif” shown in primates(*47*).

The predictive routing model(*6*, *44*) also observed θ oscillations in the course of afferent prediction-error processing, a feature found in all our oddball results but especially in the global oddball. This paper’s results suggest that ɣ synchronization routes fast transmission of feedforward stimulus information in feature-specific cortical columns(*48*) not inhibited by feedback ⍺/β synchronization; the signature of global oddballs could indicate that ⍺/β-band feedback silenced ‘Y’-responsive cortical columns, leaving ‘X’ columns to send fast ɣ-synchronized prediction-error signals, while also integrating longer-term sequence information into the slower θ-synchronized prediction error signal. This agrees with previous results (*49*) suggesting that θ oscillations may play a role in supporting sequence learning and recognition. It also fits with proposals whereby θ oscillations synchronize distant areas, enabling them to exchange relevant signals during cognitive processing(*50*, *51*). Similarly, deviance detection and release from adaptation effects showed that the peak ɣ frequency shifted up for less predictable sequences compared to more predictable sequences (Supplementary Figure 2), another potential update to the predictive routing model of predictive processing.

Where this paper found evidence for predictive routing across multiple oddball types, recent work(*17*) analyzing the same dataset in the spike-rate domain did not find evidence for predictive coding in V1 across these same oddball types. How can population-level neuronal oscillations encode prediction errors when spike rates do not? Classical experiments(*52–55*) have found that information can be more strongly encoded in the timing of individual spikes as compared to the spike rate. Computational models(*56–59*) have captured such effects, further fortifying the neurobiological plausibility of temporal coding. Newer experiments have begun to suggest that ensemble temporal codes may reveal task-modulation effects beyond those reflected in spike rates and remain robust against representational drift(*60*, *61*). Both the isocortex(*58*, *62–64*) (hippocampal-entorhinal complex) and neocortex(*65–68*) show evidence of coding information in spike times via their relationship to the phase of a circuit-level oscillation. These results together suggest a hypothetical complimentary code for representing the brain’s internal mode, with spike rates encoding stimulus representation and oscillatory synchronization encoding relative uncertainty or confidence in these representations.

Population-level modeling studies have proposed that ɣ synchronization relates to the gain of superficial pyramidal neurons(*20*, *69*, *70*); through the lens of predictive processing, ɣ oscillations may regulate feedforward prediction-error signaling(*12*, *20*, *21*). This paper’s results show general consistency with these proposals. ɣ frequency shifts have also been proposed to enhance cortical gain(*71*). Frequency shifts have been observed in the context of attention and contrast(*19*, *70*, *72*, *73*). Faster ɣ could function to have stronger downstream impact on receiving neurons(*71*). In our data, ɣ shifts to a significantly faster frequency during deviance detection (contrast DD) in V1 and LM (Supplementary Figure 2), supporting this proposal. We speculate that the deviant sequences attracted more bottom-up attention compared to the fully habituated and thereby predictable sequences. Release from stimulus-specific adaptation (contrast SSA, Supplementary Figure 2) also showed that in the sequence with higher overall uncertainty (the “yyyy” sequence in the control block with probability 𝑃 = 50%), ɣ shifted to a significantly higher peak frequency compared to the sequence with lower uncertainty (the “xxxY” in the main block with probability 𝑃 = 80%) in LM, RL, AL, and AM. We speculate that again the less probable sequences attracted more bottom-up attention compared to the fully habituated, predictable sequences early in recording, and that understanding a stimulus change with probability 𝑃 = 80% as a non-oddball may help to explain these results. Taken together, our results on enhanced ɣ power and faster ɣ frequencies are consistent with these prior theoretical proposals, suggesting that ɣ may play a role in the gain of sensory responses, although Supplementary Figure 4 shows that despite SSA and GO contrasts modulating pupil size in Westerberg et al(*74*), DD did not seem to modulate pupil size and may not have modulated synaptic gain.

Complimentary coding of stimulus content in spike rates and surprisal in spike-timing or spike-field phase coherence (as found with ɣ oscillations and prediction errors in prefrontal and anterior cingulate cortex of macaques(*75*)) would reconcile this paper’s results, and the predictive routing model, with the evidence against spike-rate coding of prediction errors in sensory cortex (*17*), although we leave measuring the relationship between the oscillations documented here and spike timing to future work. Since the local field potentials whose oscillations we analyzed here reflect neurons’ membrane potentials(*76–78*), this paper’s results remain compatible with theoretical models of predictive processing(*79–82*) in which membrane potentials, rather than spiking activity, encode the prediction-error quantity used to infer a rate-coded, evoked spike-train representation of the stimulus. This representation could consist of either samples or summary statistics of the posterior distribution while remaining compatible with existing experimental results, while spontaneous firing rates would hypothetically reflect similar representations of the prior distribution(*26*, *83*, *84*) over previously-experienced stimuli.

This paper’s results support, but also entail updating, the predictive routing model, in which surprise modulates ⍺/β oscillatory power down and ɣ oscillatory power up. These results have the potential to rescue many aspects of predictive processing hypotheses drawn from recording or neuroimaging modalities driven in significant part by local field potential oscillations, complimenting the results of Westerberg et al(*17*) showing spike-rate coding of surprise in some areas in macaque and mice with an oscillatory routing mechanism(*85*, *86*) for surprising stimuli in lower-order cortex.

## Materials and Methods

### Animals

All experiments were performed in 16 SSTAi32 and PVAi32 mice (*Mus musculus*) of both sexes (n_male_=6, n_female_=10), aged 95-128 days. All experiments on animals were conducted with the approval of the Allen Institute’s Institutional Animal Care and Use Committee.

### Surgical Procedures

The surgery pipeline at the Allen Institute is described in detail in previous reports (*40*, *87*). Briefly, A pre-operative injection of dexamethasone (3.2 mg kg^−1^, subcutaneously was administered 1 hr before surgery to reduce swelling and postoperative pain by decreasing inflammation. Mice were initially anesthetized with 5% isoflurane (1–3 min) and placed in a stereotaxic frame (Model 1900, Kopf). Isoflurane levels were maintained at 1.5–2.5% for the duration of the surgery. Body temperature was maintained at 37.5 °C. Carprofen was administered for pain management (5–10 mg/kg, subcutaneous), and atropine was administered to suppress bronchial secretions and regulate heart rhythm (0.02–0.05 mg/kg, subcutaneous). An incision was made to remove the skin, and the exposed skull was leveled with respect to pitch (bregma–lambda level), roll, and yaw. The headframe was placed on the skull and fixed with White C&B Metabond (Parkell). Once the Metabond was dry, the mouse was placed in a custom clamp to position the skull at a rotated angle of 20° to facilitate the creation of the craniotomy over the visual cortex. A circular piece of skull 5 mm in diameter was removed, and a durotomy was performed. The brain was covered by a 5-mm-diameter circular glass coverslip, with a 1-mm lip extending over the intact skull. The bottom of the coverslip was coated with a layer of polydimethylsiloxane (SYLGARD 184, Sigma-Aldrich) to reduce adhesion to the brain surface. The coverslip was secured to the skull with Vetbond (Patterson Veterinary). Kwik-Cast (World Precision Instruments) was added around the coverslip to further seal the implant, and Metabond bridges between the coverslip and the headframe well were created to hold the Kwik-Cast in place. At the end of the procedure, but before recovery from anesthesia, the mouse was transferred to a photodocumentation station to capture a spatially registered image of the cranial window.

### Electrophysiology Experiments

The Neuropixels data was acquired at the Allen Institute as part of the OpenScope project that allows the community to apply for observation on the Allen Brain Observatory platform (https://alleninstitute.org/division/mindscope/openscope/). The Neuropixels pipeline at the Allen Institute is described in detail in previous reports (*40*, *87*).

In advance of the electrophysiology experiment, intrinsic signal imaging was performed to identify visual area boundaries (*88*). An insertion window was designed for each mouse based on the identified visual areas. Mice were then habituated to head fixation and visual stimulation over two weeks.

On the day of recording, the cranial coverslip was removed and replaced with the insertion window containing holes aligned to six cortical visual areas. Mice were allowed to recover for 1–2 hours after the window was placed before being head-fixed in the recording rig.

Six Neuropixels probes were targeted to each of the six visual cortical areas (V1, LM, RL, AL, PM, AM). The boundaries of these areas and their retinotopic maps were obtained through intrinsic signal imaging, and insertion locations were planned to target regions responsive to the center of the LCD monitor. Probes were doused with CM-DiI (1 mM in ethanol; Thermo Fisher, V22888) for post hoc ex vivo probe localization. Each probe was mounted on a 3-axis micromanipulator (New Scale Technologies). The tip of each probe was aligned to its associated opening in the insertion window using a coordinate transformation obtained via a previous calibration procedure. XYZ manipulator coordinates were obtained via a previous calibration procedure where the tips of probes were aligned to the retinotopic centers of target visual areas using an image provided by intrinsic signal imaging. The operator then moved each probe into place with a joystick, with the probes fully retracted along the insertion axis, approximately 2.5 mm above the brain surface. The probes were manually lowered to the brain surface until spikes were visible on the electrodes closest to the tip. After the probes penetrated the brain to a depth of around 100 μm, they were inserted automatically at a rate of 200 μm/min (total of 3.5 mm or less in the brain). After the probes reached their targets, they were allowed to settle for 5–10 min.

Neuropixels data was acquired at 30 kHz (spike band, 500 Hz high-pass filter) and 2.5 kHz (LFP band, 1,000 Hz low-pass filter) using the Open Ephys GUI (*89*). Videos of the eye and body were acquired at 60 Hz. Pupil size was measured via previously described means (*40*). Briefly, a universal eye tracking model trained in DeepLabCut (*90*), a ResNET-50-based network, recognizes up to 12 tracking points each around the perimeter of the eye, the pupil, and the corneal reflection. The angular velocity of the running wheel was recorded at the time of each stimulus frame, at approximately 60 Hz. All LFPs were packaged into Neurodata Without Borders (NWB) files (*91*) and analyzed in this paper.

### Visual Stimulation Parameters

All Neuropixels recordings were according to the standardized Neuropixels visual coding pipeline. Visual stimuli were generated using custom PsychoPy scripts and displayed using an ASUS PA248Q LCD monitor with 1920x1200 pixels (55.7 cm wide, 60 Hz refresh rate). Stimuli were presented monocularly, and the monitor was positioned 15 cm from the right eye of the mouse and spanned 120 dva × 95 dva. Monitor placement was standardized across rigs such that the mouse’s right eye gaze would typically be directed at the center of the monitor. Stimuli consisted of full-screen oriented drifting bar gratings, oriented 45° off the vertical (and horizontal) meridian (i.e., 45° and 135° CCW from 3 o’clock). x/X refers to one orientation and y/Y, the other orientation. Sequences consisted of 4 stimuli presented for 500 ms each with a 500 ms intervening ISI and 1 s between sequences.

### Habituation to Predicted Sequence

Mice were habituated over the course of 5 behavior-only sessions in which only the predicted sequence (either xxxY or yyyX) was presented. This resulted in 2000 habituation trials before recordings.

### Data Analysis

No units from 1 mouse passed the criteria due to a destabilizing event during the recording. Additionally, data from another mouse were omitted from the analysis due to issues with the visual stimulus timing. As a result, our final sample consisted of LFPs from 14 mice. Prior to running our analyses, unless otherwise stated, LFPs were first epoched according to condition within each subject’s NWB file and the contiguous subset of channels histologically labelled as located in the cortex were selected. Channels were then aligned across subjects by locating the median channel within the granular layer L4 and determining the minimal distances in channels “down” into infragranular and “up” into supragranular cortex available across identically numbered probes in the same area across subjects. This channel range was used to align the LFPs for each subject to a shared channel numbering, and histological labellings were taken for the resulting cross-subject LFP data via majority vote among the per-subject probes. After alignment of the LFPs across subjects, they were appended in the trial dimension to collect within-condition, cross-subject LFP data. Finally, before calculating the spectrogram or spectrum for the LFPs, they were downsampled by 4 to obtain a single vertical line of channels.

Spectrograms were calculated by use of the <u>Syncopy</u> software package(*92*), used as a backend within <u>our own</u> <u>analysis code</u>. The FOOOF algorithm(*93*) (version 1.1.0) was used to parameterize neural power spectra and then remove the estimated aperiodic component from spectrograms. Settings for the algorithm were set as: peak width limits : (0.5, 12.0); max number of peaks: ∞; minimum peak height: 0.0; peak threshold: 2.0; and aperiodic mode: “fixed”. Power spectra were parameterized across the frequency range 2 to 90 Hz with a frequency resolution of 2 Hz. Unless otherwise noted, all statistics were performed using nonparametric, cluster-based permutation testing(*39*), implemented in the <u>MNE</u> package(*94*), of the LFP spectrograms at the multi-channel level, with two-tailed testing and significance thresholds set to ⍺ = 0.01.

Current source density (CSD) results came from calculating the evoked potential of oscillatory LFPs, downsampling by four (4) channels to have a single vertical column, and then employing the Elephant(*95*) package’s CSD implementation. The oscillatory LFPs themselves came from replicating the analysis of van Kerkoerle et al(*31*) (among others(*96*, *97*)): we first band-pass the raw (though epoched) LFPs in chosen oscillatory bands (2-10 Hz for θ, 10-30 Hz for ⍺/β, and 30-90 Hz for ɣ) and obtain the oscillatory phases from Sycopy(*92*); we then selected the time and trial coordinates in the original epoched LFPs where the absolute value of the phase came within 0.001 of π (denoting an oscillatory trough) and, after detrending of the original LFPs by subtracting its trial-averaged evoked potential, re-epoched for an oscillatory period around those timepoints in those trials. Current source density was then again calculated via Elephant.

## Data and Code Availability

All mouse data used in this work are openly available through the DANDI Archive (dandiset id: 000253, doi: https://doi.org/10.48324/dandi.000253/0.240503.0152). Software resources for the analysis of this dataset are provided here: https://github.com/BastosLab/epych/tree/develop/notebooks/passiveglo. The code to generate the global-local oddball sequences is provided here: https://github.com/AllenInstitute/openscope-glo-stim.

## Acknowledgments

The Neuropixels dataset was obtained at the Allen Brain Observatory as part of the OpenScope program, which is operated by the Allen Institute, Neural Dynamics. **Funding:** This research was supported by the National Institute of Mental Health (NIMH) R00MH116100, Vanderbilt University startup funds, a Vanderbilt Brain Institute Faculty Fellow Award, the NARSAD Young Investigator Award from the Brain and Behavior Research Foundation, and the National Science Foundation (NSF) Faculty Early Career Development Program (CAREER) grant 2339210. J.A.W. was supported by a fellowship from the International Human Frontier Science Program Organization (HFSPO) LT0001/2023-L and a grant from the Dutch Research Council VI.Veni.232.110.

## Author contributions

Conceptualization: ES, JAW, AMB

Data curation: ES, JAW

Conceptualization: ES, JAW, AMB

Data curation: ES, JAW

Formal analysis: ES

Funding acquisition: JAW, AMB

Supervision: JSS, AMB

Visualization: ES

Writing – original draft: ES, AMB

Writing – review, and editing: all authors

## Supplementary Materials

**Supplementary Figure 1:**
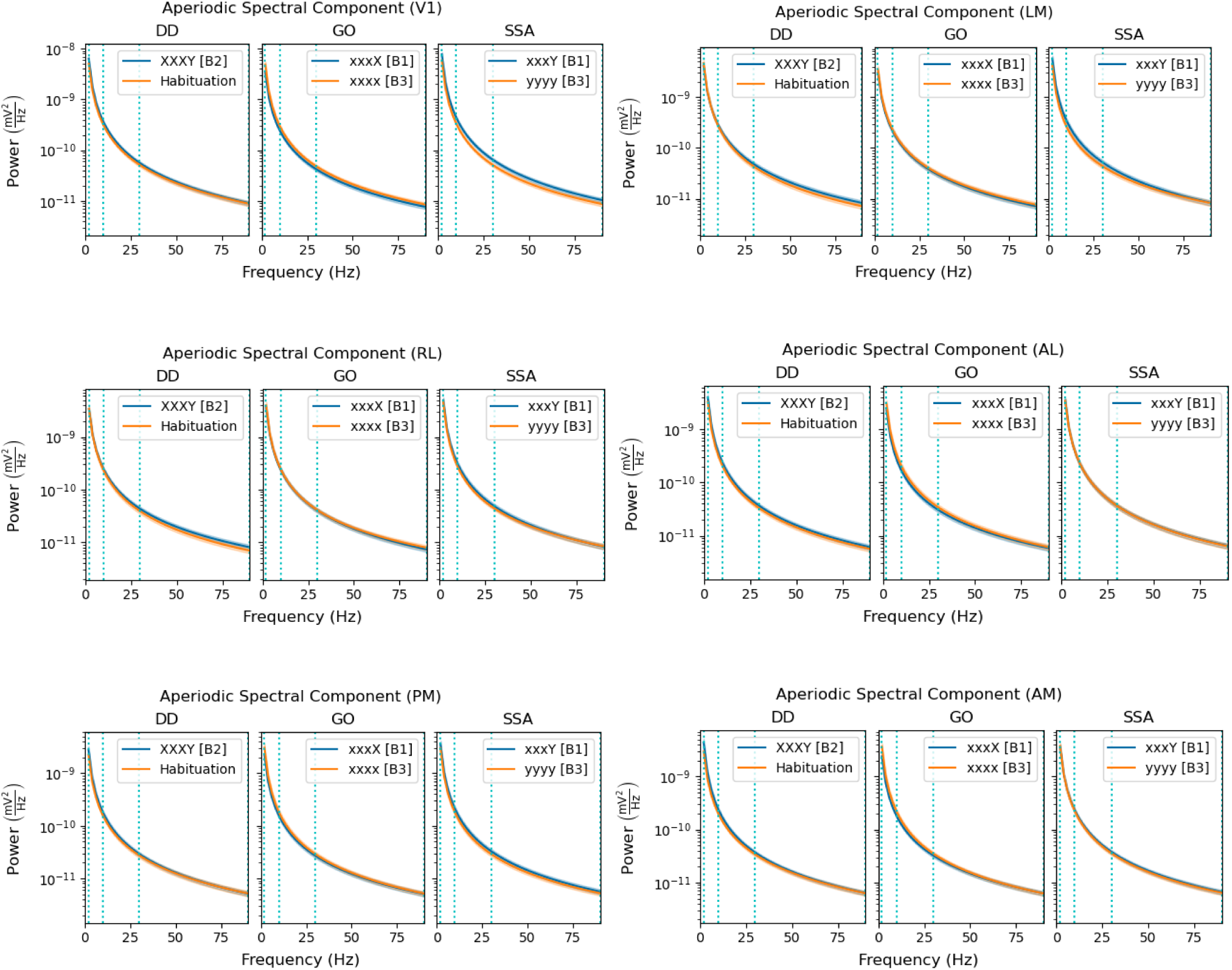
Aperiodic components of spectral responses lack significant cross-condition differences. Fitting parametric models using FOOOF (*93*)to the trial-averaged, within-subject (N=14) spectral responses to oddballs in areas V1, LM, RL, AL, PM, and AM separated out the aperiodic component (power shown in mV^2^/Hz in log-scale) from the broader spectral response; the six plots here each show this aperiodic component for the oddball (blue) and control (yellow) stimuli/sequences across the three contrasts, including error bands of two standard errors of the mean (SEMs). Across all three contrasts, the error bands for the oddball and control stimuli overlap closely enough to lack statistical significance. The blue, vertical dotted lines show the boundaries between θ (2-10 Hz), ⍺/β (10-30 Hz), and ɣ (30-90 Hz) frequency bands.

**Supplementary Figure 2:**
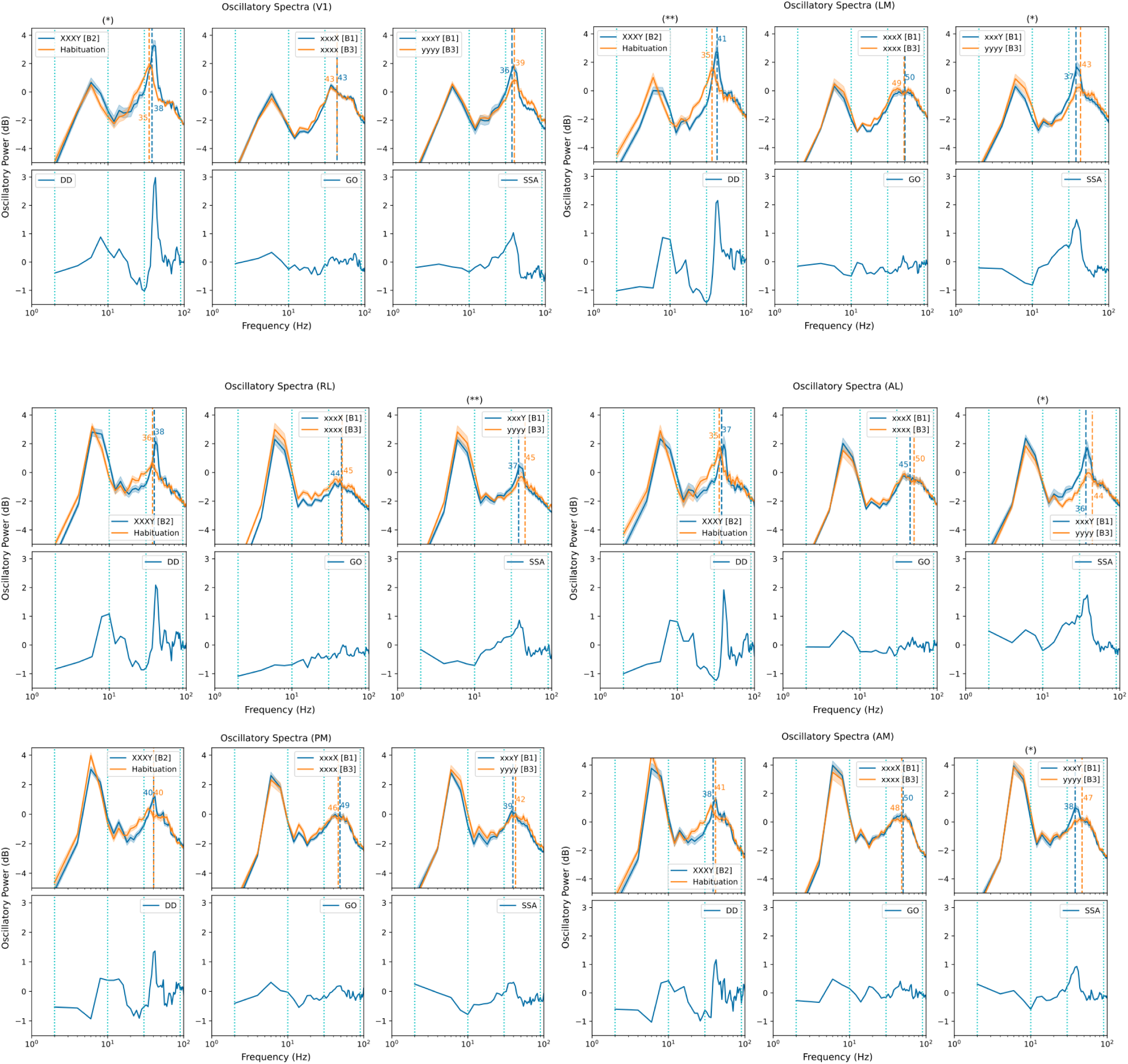
Oscillatory spectra show significant differences in power of ⍺/β and ɣ peaks in deviance detection and stimulus-specific adaptation contrasts across areas, while global oddball contrasts show limited significant differences. After removing the aperiodic components shown in Supplementary Figure 1, the oscillatory peaks are shown here in log-space (decibels) across areas (V1, LM, RL, AL, PM, and AM) and contrasts (DD, GO, and SSA), with error-bands of two standard errors of the mean (SEMs) shown surrounding each average spectrum. DD and SSA consistently show significant (>2 SEM) differences in the heights (power above aperiodic baseline) of the ɣ (30-90 Hz) oscillatory peaks in V1, LM, RL, AL, and AM, with DD showing such a significant peak in PM as well. DD shows significant difference in heights of the ⍺/β (10-30 Hz) peaks, with the control condition showing greater ⍺/β power than the oddball condition, in V1, LM, RL, AL, PM, and AM; in LM and AL the SSA contrast shows higher ⍺/β power in the oddball condition rather than the control condition. In V1, LM, RL, AL, and PM the GO contrast shows higher ⍺/β power in the control condition. In DD, the oddball’s ɣ-band peak frequency significantly (in a permutation-based *t*-test with 1000 permutations) shifted higher than that of the control condition V1 (35-38 Hz, ⍺ < 0. 05) and LM (35-41 Hz, ⍺ < 0. 01). In SSA, the “oddball” (having probability 𝑃 = 80%) had a ɣ-band peak frequency shifted significantly (in a permutation-based *t*-test with 1000 permutations) to a *lower* frequency in LM (37-43 Hz, ⍺ < 0. 05), RL (37-45 Hz, ⍺ < 0. 01), AL (36-44 Hz, ⍺ < 0. 05), and AM (38-47 Hz, ⍺ < 0. 05). Statistics here were calculated within conditions and across subjects, averaging over channels and trials.

**Supplementary Figure 3:**
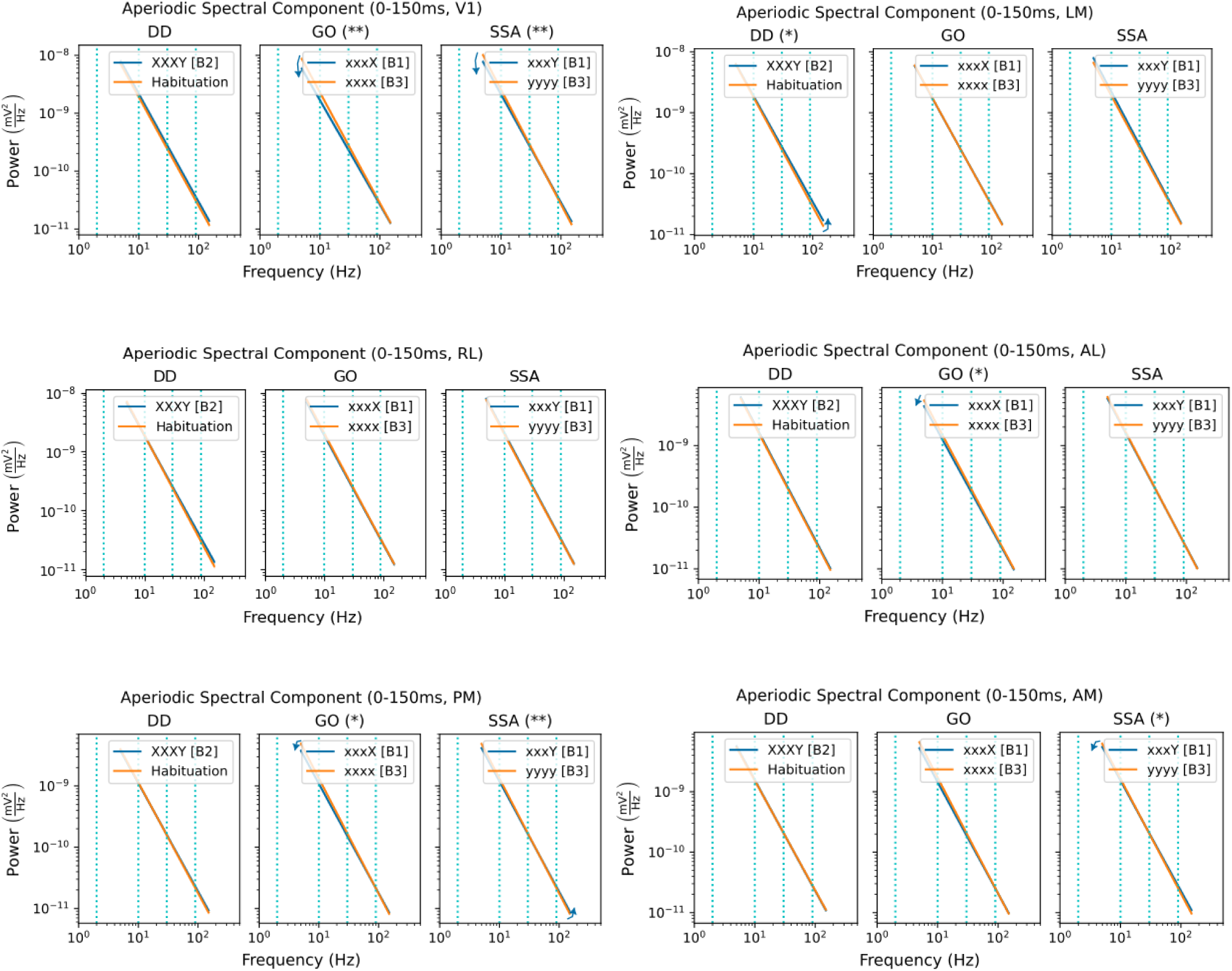
Aperiodic components of spectral responses in the first 150ms after oddball onset display significant cross-condition differences. Fitting parametric models using FOOOF(*93*) to the trial-averaged, within-subject (N=14) first 150 ms of the spectral responses to oddballs in areas V1, LM, RL, AL, PM, and AM separated out the aperiodic component (power shown in mV^2^/Hz in log-scale) from the broader spectral response; the six plots here each show this aperiodic component for the oddball (blue) and control (yellow) stimuli/sequences across the three contrasts, including error bands of two standard errors of the mean (SEMs). Contrasts and areas marked with an asterisk (DD-LM, GO-AL, GO-PM, SSA-AM) displayed at least one cross-condition difference in the two parameters of the aperiodic component with significance level ⍺ < 5%, while those marked with two asterisks (GO-V1, SSA-V1, SSA-PM) displayed at least one cross-condition difference with significance level ⍺ < 1%; significance was measured by a permutation-based t-test on the two independent coordinates with 1000 permutations each. The blue, vertical dotted lines show the boundaries between θ (2-10 Hz), ⍺/β (10-30 Hz), and ɣ (30-90 Hz) frequency bands.

**Supplementary Figure 4:**
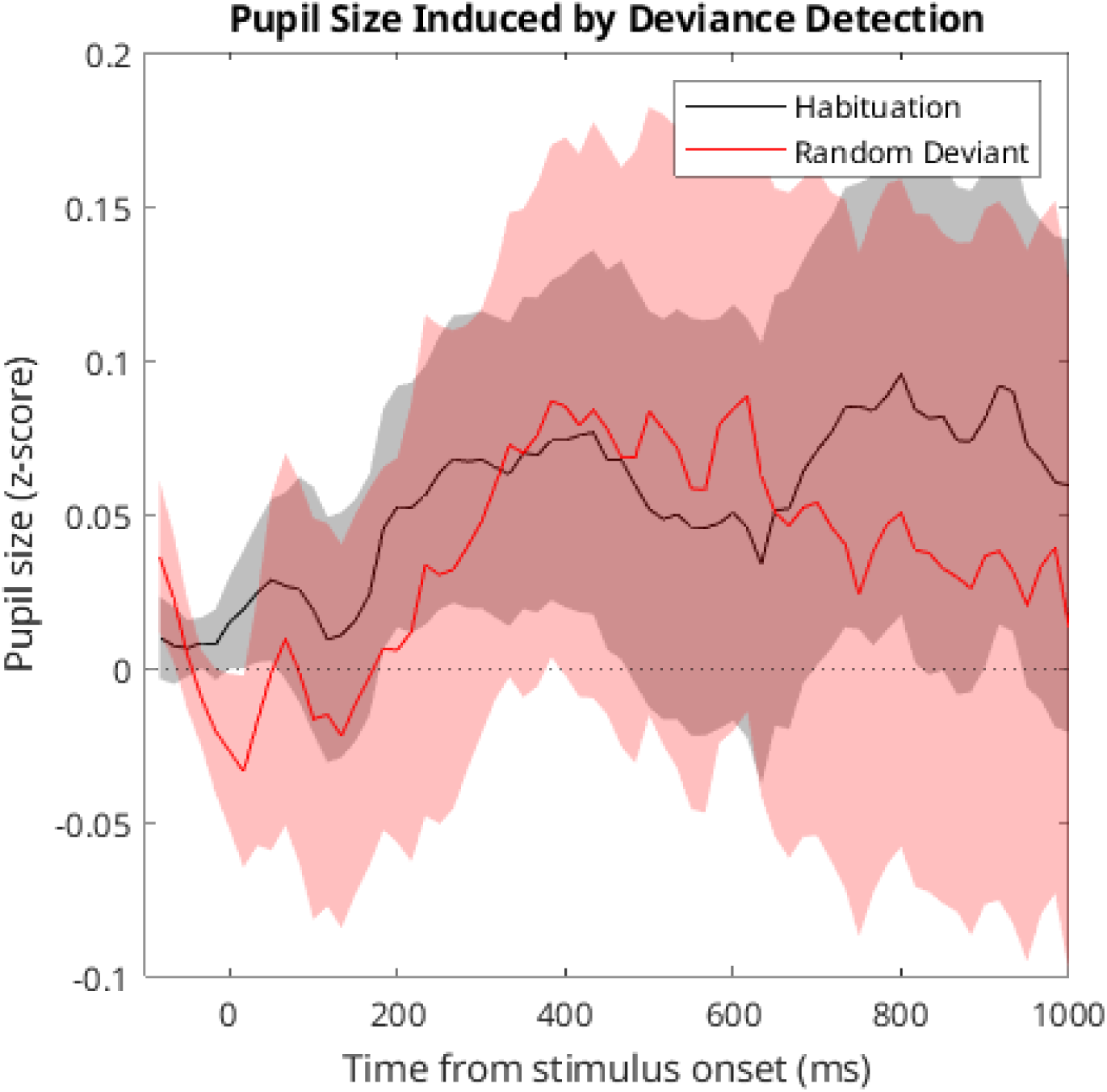
Deviance detection does not significantly change pupil size. Taking the z-scores of pupil sizes across P4 of the “XXXY” random control sequences and “xxxy” habituation sequences did not show significant differences. Solid lines show the mean z-score of pupil size across trials, while error bands show two trialwise standard deviations of the mean (trialwise standard deviation over square-root of number of trials). The dotted line shows the mean pupil size overall with a z-score of zero.

